# *XRN1* deletion induces PKR-dependent cell lethality in interferon-activated cancer cells

**DOI:** 10.1101/2023.08.01.551488

**Authors:** Tao Zou, Meng Zhou, Akansha Gupta, Patrick Zhuang, Alyssa R. Fishbein, Hope Y. Wei, Zhouwei Zhang, Andrew D. Cherniack, Matthew Meyerson

**Affiliations:** Department of Medical Oncology, Dana-Farber Cancer Institute, Boston, Massachusetts 02215, USA; Broad Institute of Harvard and MIT, Cambridge, Massachusetts 02142, USA; Center for Cancer Genomics, Dana-Farber Cancer Institute, Boston, Massachusetts 02215, USA; Department of Genetics, Harvard Medical School, Boston, Massachusetts 02115, USA

**Keywords:** XRN1, PKR, RNA sensing, interferon, cancer

## Abstract

Emerging data suggest that induction of viral mimicry responses through activation of double-stranded RNA (dsRNA) sensors in cancer cells is a promising therapeutic strategy. One approach to induce viral mimicry is to target molecular regulators of dsRNA sensing pathways. Here, we show that the exoribonuclease XRN1 is a negative regulator of the dsRNA sensor protein kinase R (PKR) in cancer cells with high interferon-stimulated gene (ISG) expression. *XRN1* deletion causes PKR activation and consequent cancer cell lethality. Disruption of interferon signaling with the JAK1/2 inhibitor ruxolitinib can decrease cellular PKR levels and rescue sensitivity to *XRN1* deletion. Conversely, interferon-β stimulation can increase PKR levels and induce sensitivity to *XRN1* inactivation. Lastly, *XRN1* deletion causes accumulation of endogenous complementary sense/anti-sense RNAs, which may represent candidate PKR ligands. Our data demonstrate how XRN1 regulates PKR and nominate XRN1 as a potential therapeutic target in cancer cells with an activated interferon cell state.

## Introduction

Anti-tumor immune responses rely on the ability of the immune system to recognize tumor cells as non-self^1^. For example, adaptive immune cells can recognize cancer neoantigens as non-self to initiate tumor cell killing^2^. This mechanism of self/non-self recognition underlies the efficacy of immune checkpoint inhibitors and adoptive cellular therapies, which have improved clinical outcomes for patients with hematologic and solid tumor malignancies^1, 2^. Similar to the adaptive immune system, the innate immune system also possesses mechanisms to distinguish self from non-self^3^, but it is unclear whether these pathways can be leveraged for cancer therapeutics.

Recent studies have suggested that specific genetic or pharmacologic perturbations can induce the expression of endogenous immunogenic RNAs that can activate innate immune dsRNA sensing pathways in cancer cells^4–15^, a process termed as “viral mimicry^4, 14, 15^.” Engagement of these RNA sensing pathways can, in turn, cause direct cancer cell cytotoxicity and/or stimulate anti-tumor adaptive immune responses. Disruption of epigenetic regulation^4–6, 10^, perturbation of RNA splicing^11^, and inhibition of protein arginine methylation^13^ can all trigger the expression of dsRNA, including endogenous retroelements, and stimulate dsRNA sensing and interferon responses to inhibit cancer cell growth.

In parallel with these studies, our group and others have shown that depletion of the adenosine deaminase of RNAs (ADAR1) causes activation of the dsRNA sensor PKR in cancer cell lines with high levels of ISG expression^7, 8, 12^. PKR is encoded by the *EIF2AK2* gene and normally detects dsRNA of viral origin. Binding of viral dsRNA ligands triggers PKR dimerization and autophosphorylation, activating its kinase function to initiate signals that ultimately inhibit cellular protein translation, thereby restricting viral replication^16, 17^. Because PKR detects dsRNA in a sequence agnostic manner, endogenous cellular dsRNA can also stimulate its activation^16, 18^. To this point, a subsequent study showed that DNA hypomethylating agents can induce a requirement for ADAR1 in cancer cells through the induction of inverted *Alu* elements^19^, which, together with mitochondrial RNA^20^, are known activating ligands of PKR. However, the range of cellular processes that regulate PKR activation and the identity of endogenous RNAs capable of stimulating PKR remain incompletely defined.

We hypothesized that cancer cell lines displaying a requirement for ADAR1 may also require other molecular regulators of dsRNA sensors for their survival. In this study, we analyzed genome-scale CRISPR-Cas9 gene essentiality screening data across hundreds of cancer cell lines and identified a correlation between the requirement for ADAR1 and the XRN1 exoribonuclease in cancer cell lines with high levels of ISG expression. Mechanistically, *XRN1* deletion activates PKR, which was required for cancer cell lethality after XRN1 depletion. To understand the role of interferon pathway activation, we showed that inhibition of JAK1/2 signaling can decrease cellular PKR levels and rescue *XRN1* KO-sensitivity. In conjunction, stimulation of interferon signaling in *XRN1* KO-insensitive cell lines increased PKR levels and induced sensitivity to *XRN1* deletion in a PKR-dependent manner. Computational analysis of RNA sequences revealed an accumulation of complementary sense/anti-sense RNA transcripts after *XRN1* deletion. We propose that *XRN1* deletion in cancer cell lines with high levels of the interferon-stimulated dsRNA sensor PKR causes the accumulation of complementary sense/anti-sense RNAs pairs, thereby causing PKR activation and cancer cell lethality.

## Results

### The enzymatic activity of XRN1 is required for survival of cancer cell lines with high expression of ISGs

To uncover molecular regulators of dsRNA sensors that may function similarly to ADAR1, we analyzed genome-scale CRISPR-Cas9 screening data generated by the Cancer Dependency Map^21^. *XRN1* genetic dependency was the top correlate with *ADAR1* genetic dependency across cancer cell lines, followed by *PRKRA* and *USP18* (**Figure 1A**). Notably, this correlation between *ADAR1* and *XRN1* dependency held across multiple cancer lineages and degrees of gene essentiality, with an R^2^ of ∼0.48 (**Figure 1B**). *XRN1* encodes an exoribonuclease that functions as the major pathway for cytoplasmic 5’ to 3’ RNA decay^22^. *XRN1* gene essentiality in cancer cell lines is correlated with the requirement for multiple genes beyond *ADAR1*, several of which encode proteins involved in RNA degradation and metabolism, including the RNA de-capping proteins DCP2 and EDC4 (**Figure 1C**). Similar to *ADAR1* gene essentiality, cancer cell lines that require *XRN1* displayed an enrichment in gene sets that represent ISG expression (**Figure 1D**). To validate the results of this analysis, we used CRISPR-Cas9 to target control genes or *XRN1* (Figure S1A) in cancer cell lines. We found that lung cancer cell lines that require *ADAR1* showed a significant cell viability defect after *XRN1* deletion compared to control cells, whereas lung cancer cell lines that do not require *ADAR1* also did not require *XRN1* for their survival (**Figure 1E**). Next, we selected cancer cell lines from diverse lineages, including pancreatic adenocarcinoma, mesothelioma, triple negative breast carcinoma, melanoma, and gastric adenocarcinoma, that were predicted to require both *ADAR1* and *XRN1*. *XRN1* deletion in each of these cell lines caused a cell viability defect compared to controls (**Figure 1F**). Taken together, these data show that cancer cell lines that require *ADAR1* also require *XRN1* for survival, suggesting that these genes may function through similar pathways to control cancer cell viability.

**Figure 1.**
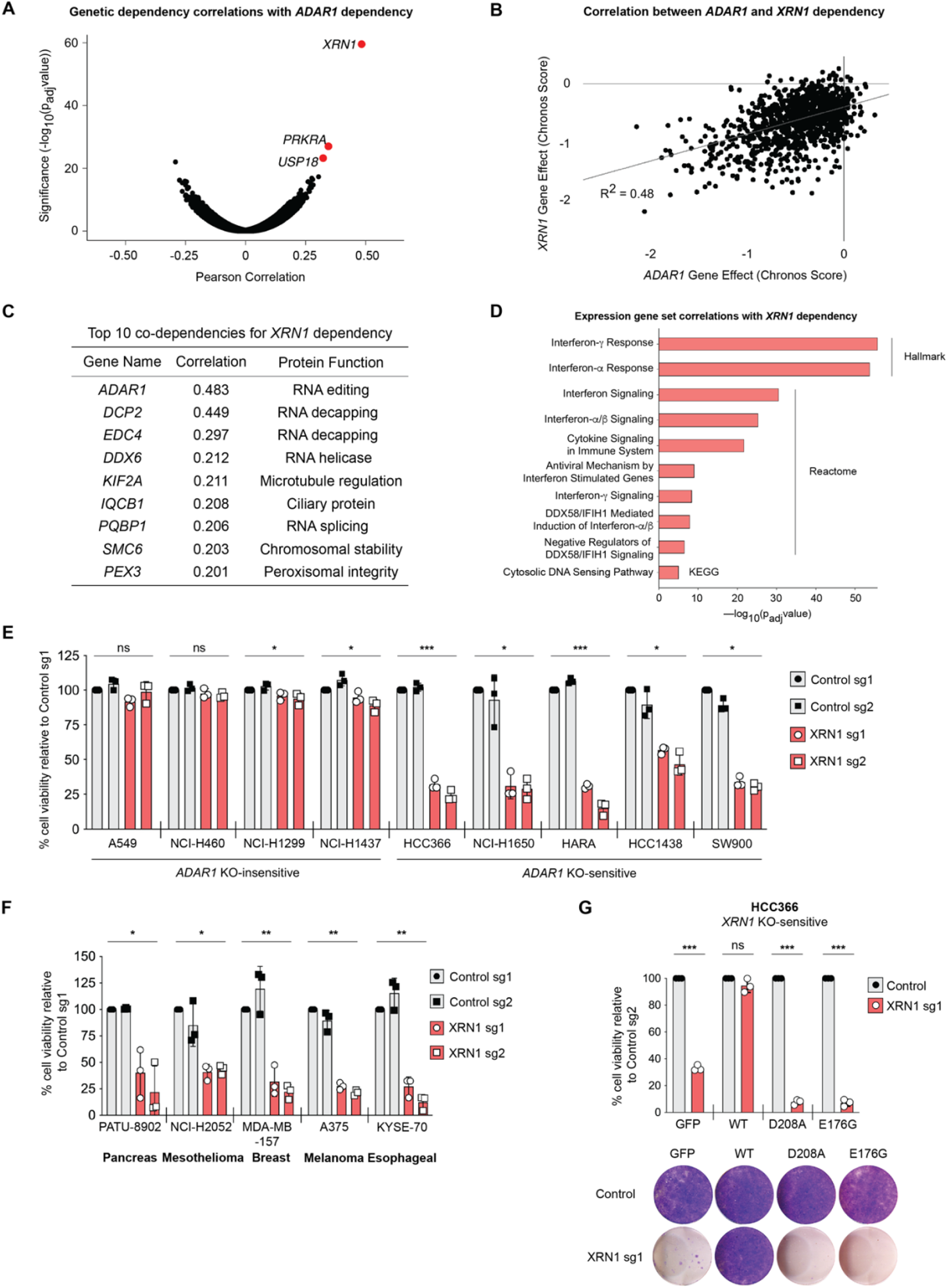
The enzymatic activity of XRN1 is required for survival of cancer cell lines with high expression of ISGs. (**A**) Correlation and statistical significance of genome-wide genetic dependencies compared to *ADAR1* genetic dependency based on CRISPR-Cas9-mediated gene essentiality screens. Each dot represents a different gene. Pearson correlations and corresponding adjusted *p* (*p*_adj_) values were computed for each feature in the Cancer Dependency Map Public 22Q4 dataset using all cancer cell lines. (**B**) Correlation of *ADAR1* and *XRN1* genetic dependency based on CRISPR-Cas9-mediated gene essentiality screens. CHRONOS^43^ is a computational method that predicts the sensitivity of cell lines to deletion of specific genes using data from gene essentiality screens. Lower CHRONOS scores predict for cell lines that are relatively sensitive to gene deletion, while higher scores predict for cell lines that are relatively insensitive to gene deletion. Each dot represents a different human cancer cell line. Pearson correlations were computed for each feature in the Cancer Dependency Map Public 22Q4 dataset using all cancer cell lines. (**C**) Top 10 genetic dependencies correlated with *XRN1* genetic dependency in CRISPR-Cas9-mediated gene essentiality screens. Pearson correlations were computed for each feature in the Cancer Dependency Map Public 22Q4 dataset using all cancer cell lines. (**D**) Expression gene set correlations with *XRN1* genetic dependency based on CRISPR-Cas9-mediated gene essentiality screens. Pearson correlations and corresponding *p*_adj_ values were computed for each feature in the Cancer Dependency Map Public 22Q4 dataset using all cancer cell lines. A subset of the 200 genes with the highest absolute correlation was used to calculate gene set over-representation. C2 Canonical Pathways v7.0 and Hallmark v7.1 gene set collections from MSigDB were analyzed. *p* values were computed using a hypergeometric test and then corrected for multiple hypothesis testing using Benjamini/Hochberg false discovery rate correction to generate *p*_adj_ values. (**E**) Lung cancer cell viability was assessed by ATP bioluminescence 12 days after targeting control loci (sg1: AAVS, sg2: *Chr2.2*) or *XRN1* (sg1: exon 6, sg2: exon 19) with CRISPR-Cas9. (**F**) Cell viability was assessed by ATP bioluminescence 12 days after targeting control loci or *XRN1* with CRISPR-Cas9 in cancer cell lines of diverse lineages. (**G**) Cell viability was assessed by ATP bioluminescence (top panel) or crystal violet staining (bottom panel) 12 or 17 days, respectively, after targeting of a control locus or endogenous *XRN1,* by XRN1 sg1, with CRISPR-Cas9 in HCC366 cells expressing *GFP* control or overexpressing WT *XRN1* resistant to *XRN1* sg1 targeting or catalytically inactive mutants of *XRN1* (D208A and E176G) resistant to *XRN1* sg1 targeting. Raw data from gene essentiality screens in (**A**-**D**) were obtained from the Cancer Dependency Map^21^. ATP bioluminescence values were normalized to the control sg1 sample within each cell line. Data from three independent biological replicates are shown in (**E**-**G**). Error bars represent standard deviation. \**p* < 0.05, \*\**p* < 0.01, *** *p* < 0.001, and ns = not significant, as calculated by repeated measures one-way analysis of variance (ANOVA) in (**E**) and (**F**) or paired Student’s *t* test in (**G**).

To determine whether the effects of XRN1 depletion are on-target, we generated wildtype *XRN1* constructs that are resistant to cleavage by *XRN1* guides sg1 or sg2 and overexpressed these constructs in *XRN1* knockout (KO)-sensitive HCC366 and NCI-H1650 cells (Figures S1B-D). Overexpression of sg1/sg2-resistant wildtype *XRN1* rescued deletion of endogenous *XRN1* (**Figure 1G**; Figures S1E-G). To test whether the enzymatic function of XRN1 is necessary for the survival of *XRN1* KO-sensitive cancer cells, we generated sg1/sg2-resistant mutant versions of human *XRN1* (D208A and E176G) that are orthologous to XRN1-inactivating mutations in *K. lactis*^23^ and *D. melanogaster*^24^. Overexpression of *XRN1* D208A and E176G failed to rescue cell viability in *XRN1* KO-sensitive cancer cells after deletion of endogenous *XRN1* (**Figure 1G**; Figures S1E-G), demonstrating a requirement for XRN1 catalytic activity. Of note, HCC366 cells expressing the catalytically inactive versions of XRN1 (D208A and E176G) appear to have decreased cell viability compared to the GFP-expressing controls cells after endogenous *XRN1* deletion (**Figure 1G**; Figure S1E), suggesting a possible dominant negative function for these XRN1 mutations. However, we did not observe a similar effect in NCI-H1650 cells (**Figures S1F** and **S1G**) and HCC366 and NCI-H1650 expressing control sgRNAs did not show a substantial growth defect (**Figure 1G**; Figures S1E-G). Together, these data indicate that the enzymatic function of XRN1 is necessary for cancer cell survival in the setting of high ISG expression.

### PKR is required for cancer cell lethality after *XRN1* deletion

Given prior work showed that PKR contributes to *ADAR1* genetic dependency in cancer cells^7, 8, 12^ and that viral infection of XRN1-deficient cells activates PKR^25, 26^, we examined whether PKR activation contributes to cancer cell lethality after *XRN1* deletion. While *XRN1* deletion did not increase PKR phosphorylation in *XRN1* KO-insensitive cell lines, including A549 and NCI-H460 (**Figure 2A**), as well as NCI-H1299 and NCI-H1437 (**Figure S2A**), XRN1 loss led to increased PKR phosphorylation specifically in *XRN1* KO-sensitive cell lines, including HCC366 and NCI-H1650 (**Figure 2A**), as well as HARA and HCC1438 (**Figure S2A**). RNA-sequencing showed that *XRN1* deletion in KO-sensitive cells resulted in increased expression of genes involved in the integrated stress response, NF-κB signaling, and interferon response pathways (**Figures 2B; Figure S2B; Table S1**), all of which are known downstream targets of PKR signaling^17^. Moreover, *XRN1* deletion in KO-sensitive cell lines caused increased stress granule formation as measured by G3BP1 re-localization, a marker of PKR activation^27, 28^, compared to control cells (**Figure 2C**). Next, we interrogated the requirement of PKR and other dsRNA sensing pathways for *XRN1* genetic dependency. Co-deletion of *PKR* with *XRN1* deletion rescued cell viability in KO-sensitive cells (**Figure 2D; Figure S2C**) and largely reversed the gene expression changes induced by *XRN1* deletion alone (**Figure S2D; Table S2**). In contrast, deletion of *MAVS* or *RNase L* with *XRN1* deletion had no effect on cancer cell viability (**Figures 2E-F; Figures S2E-S2F**). These data demonstrate that *XRN1* deletion causes activation of the dsRNA sensor PKR, and that PKR is required for *XRN1* KO-sensitivity.

**Figure 2.**
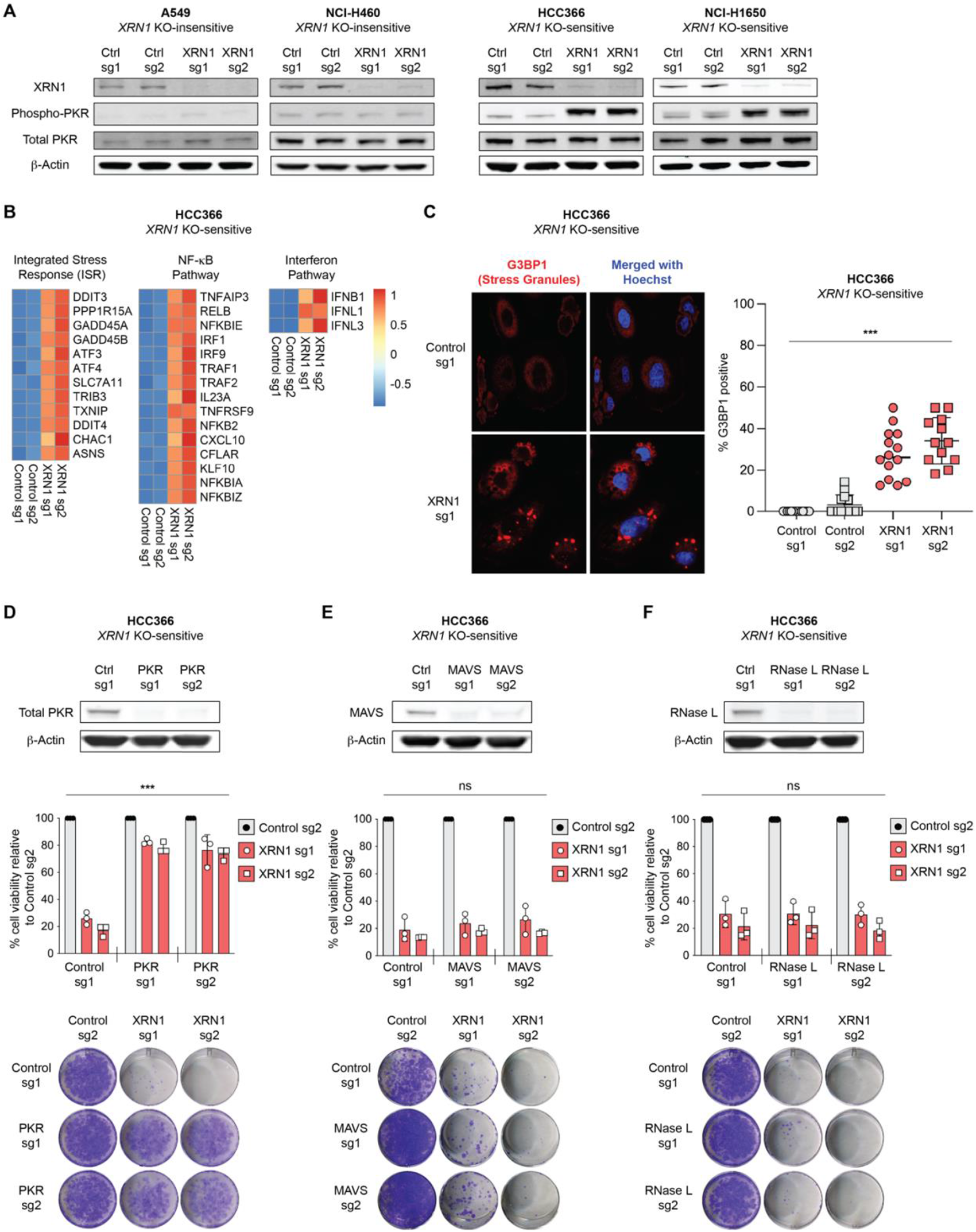
PKR signaling is required for *XRN1* gene essentiality. (**A**) Representative immunoblots showing XRN1, phospho-PKR, total PKR, and β-actin protein levels in *XRN1* KO-insensitive (A549 and NCI-H460) and *XRN1* KO-sensitive (HCC366 and NCI-H1650) cells 7 days after CRISPR-Cas9 targeting of control loci or *XRN1*. At least three independent biological replicates were performed for each cell line. (**B**) Heatmaps showing relative transcripts per million (TPM) values for the indicated differentially expressed genes in the integrated stress response, NF-κB, and interferon pathways (rows) 7 days after CRISPR-Cas9 targeting of control loci or *XRN1* (columns) in HCC366 cells. **RED**: increased expression, **BLUE**: decreased expression, as shown on the right side of the diagram. Each condition includes at least three independent biological replicates. (**C**) Immunofluorescence analysis of **G3BP1** and **Hoechst** staining 7 days after CRISPR-Cas9 targeting of control loci or *XRN1* in HCC366 cells (left panels). The percentage of cells containing at least one focus of G3BP1 staining was calculated in multiple fields across two independent biological experiments (right panel). (**D-F**) (Top panels) Immunoblots confirming depletion of PKR (**D**), MAVS (**E**), or RNase L (**F**) protein levels after CRISPR-Cas9 targeting in HCC366 cells. β-actin was used as a loading control. (Middle panels) Cell viability was assessed by ATP bioluminescence 12 days after CRISPR-Cas9 targeting of control loci or *XRN1* in PKR-depleted (**D**), MAVS-depleted (**E**), or RNase L-depleted (**F**) HCC366 cells. ATP bioluminescence values were normalized to the control sg1 sample within each cell line. Data from three independent biological replicates are shown. Error bars represent standard deviation. *** *p* < 0.001 and ns = not significant, as calculated by repeated measures two-way ANOVA. (Bottom panels) Crystal violet staining 17 days after CRISPR-Cas9 targeting of control genes or *XRN1* in PKR-depleted (**D**), MAVS-depleted (**E**), or RNase L-depleted (**F**) HCC366 cells. Crystal violet images are representative of three independent biological experiments.

### Inhibition of interferon signaling with the JAK inhibitor ruxolitinib can decrease PKR levels and rescue sensitivity to XRN1 loss

Given that the interferon-inducible protein PKR is required for *XRN1* KO-sensitivity, we reasoned that there may be a mechanistic link between elevated ISG expression and *XRN1* genetic dependency. Indeed, the most significant gene expression correlates with *XRN1* KO-sensitivity across cancer cell lines are ISGs, as shown in **Figure 1D** above, including *PKR* as the 5^th^ most correlated gene to *XRN1* KO-sensitivity (**Figure 3A**). Immunoblotting also confirmed that PKR protein levels are slightly higher (average of 1.92-fold, *p*=0.0025) in *XRN1* KO-sensitive cell lines compared to KO-insensitive cell lines (**Figure S3A**). Thus, we hypothesized that activated interferon signaling may increase cellular PKR levels, thereby sensitizing cancer cell lines with high ISG expression to *XRN1* deletion.

**Figure 3.**
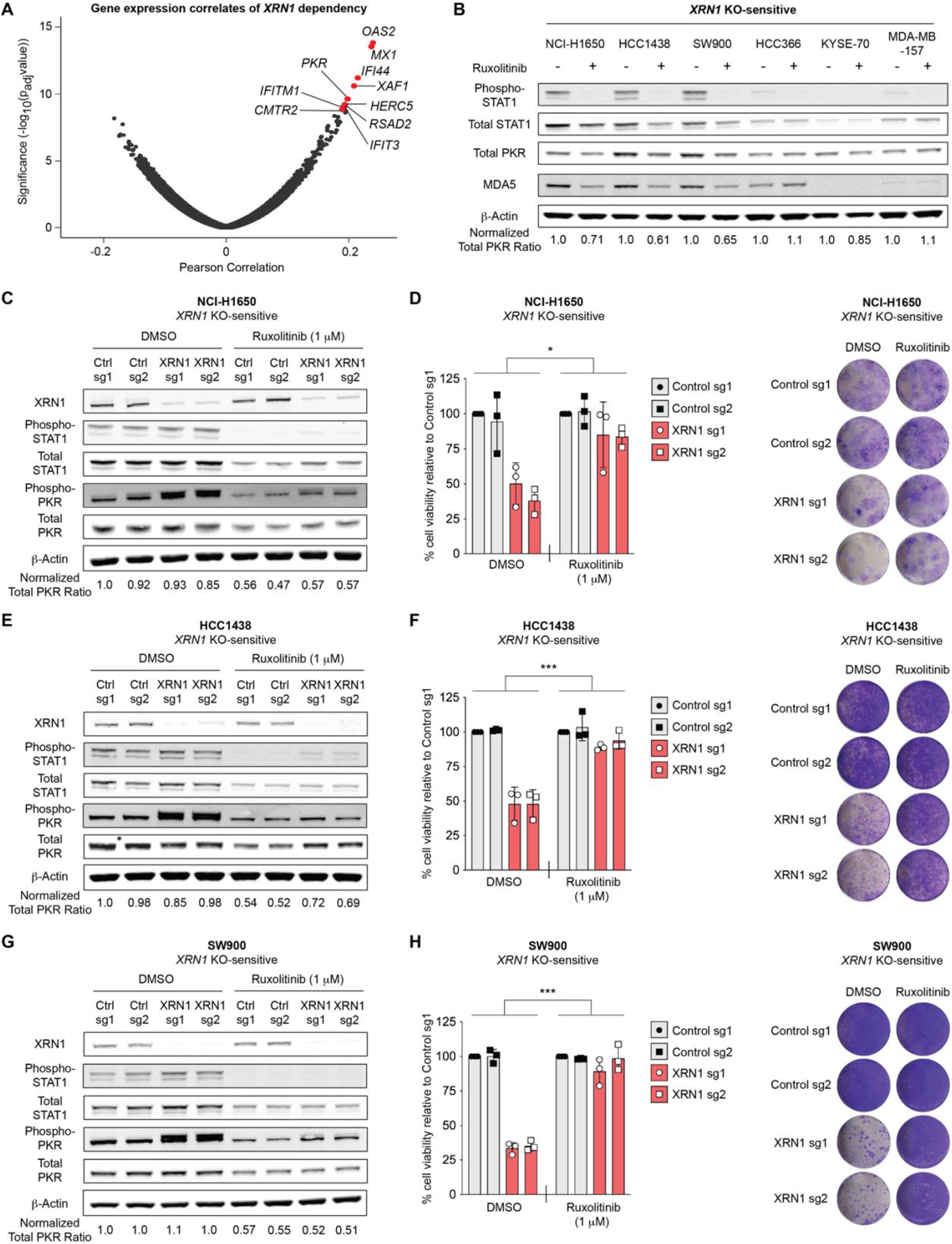
Inhibition of JAK1/2 signaling can modulate PKR levels and sensitivity to *XRN1* deletion. (**A**) Correlation of expression levels of individual genes with *XRN1* genetic dependency based on CRISPR-Cas9-mediated gene essentiality screens. Each dot represents one gene and the top 10 gene expression correlates are labeled in **RED**. Pearson correlations and corresponding *p*_adj_ values were computed for each feature in the Cancer Dependency Map Public 22Q4 dataset using all cancer cell lines. (**B**) Representative immunoblots showing phospho-STAT1, total STAT1, total PKR, MDA5, and β-actin protein levels in a panel of *XRN1* KO-sensitive cancer cell lines treated with either DMSO control or ruxolitinib (1μM) for 24 hours. (**C, E, G**) Representative immunoblots showing XRN1, phospho-STAT1, total STAT1, phospho-PKR, total PKR, and β-actin protein levels in control and *XRN1*-deleted NCI-H1650 (**C**), HCC1438 (**E**), and SW900 (**G**) cells treated with either DMSO control or ruxolitinib (1μM). Three independent biological replicates were performed for each cell line. (**D**, **F**, **H**) Cell viability was assessed by either ATP bioluminescence (left panels) or crystal violet staining (right panels) after CRISPR-Cas9 targeting of control loci or *XRN1* in NCI-H1650 (**D**), HCC1438 (**F**), and SW900 (**H**) cells treated with DMSO control or ruxolitinib (1μM). ATP bioluminescence values were normalized to the control sg1 sample within each cell line. Data from three independent biological replicates are shown. Error bars represent standard deviation. \**p* < 0.05 and *** *p* < 0.001, as calculated by repeated measures two-way ANOVA. Crystal violet images are representative of three independent biological experiments.

To interfere with cellular interferon signaling, we utilized the selective JAK1/2 inhibitor ruxolitinib^29^, which caused a dose-dependent inhibition of STAT1 phosphorylation and downregulation of ISG protein levels in cancer cell lines with high basal activation of the interferon pathway (**Figure 3B; Figure S3B**). Of note, a subset of *XRN1* KO-sensitive cancer cell lines displayed higher basal levels of STAT1 phosphorylation (**Figure 3B**).

To determine the contribution of JAK1/2-dependent interferon signaling to *XRN1* KO-sensitivity, we treated control and *XRN1* KO cells with vehicle (DMSO) or ruxolitinib. Ruxolitinib treatment decreased total PKR levels and attenuated PKR phosphorylation after *XRN1* deletion as compared to DMSO treatment specifically in a group of *XRN1* KO-sensitive cell lines, namely NCI-H1650, HCC1438, and SW900 (**Figures 3C, 3E, 3G**), with higher levels of basal STAT1 phosphorylation (**Figure 3B**). In conjunction with decreases in total and phospho-PKR levels, ruxolitinib treatment largely rescued cell viability after *XRN1* deletion in NCI-H1650, HCC1438, and SW900 cells (**Figures 3D, 3F, 3H**).

Conversely, in *XRN1* KO-sensitive cell lines with lower levels of basal STAT1 phosphorylation, specifically HCC366, KYSE-70, and MDA-MB-157 cells (**Figure 3B**), ruxolitinib treatment did not decrease total PKR levels and only attenuated PKR phosphorylation modestly after *XRN1* deletion as compared to DMSO treatment (**Figures S3C, S3E, S3G**). Consequently, ruxolitinib treatment did not rescue cell viability after *XRN1* deletion in HCC366, KYSE-70, or MDA-MB-157 cells (**Figures S3D, S3F, S3H**). These data suggest that for a subset of *XRN1* KO-sensitive cancer cell lines with elevated baseline STAT1 phosphorylation, the lethality upon *XRN1* deletion is rescued by disruption of interferon signaling via inhibition of JAK1/2.

### Activation of interferon signaling can increase PKR levels and sensitivity to XRN1 loss

To extend the concept that interferon pathway activation can modulate *XRN1* KO-sensitivity, we tested whether stimulation of interferon signaling can induce vulnerability to XRN1 depletion in *XRN1* KO-insensitive cells with low levels of basal ISG expression, similar to our previous report that interferon stimulation sensitizes cancer cells to ADAR1 depletion^7^. Interferon-β stimulation of *XRN1* KO-insensitive cell lines (A549, NCI-H460, NCI-H1299, and NCI-H1437) increased levels of STAT1 phosphorylation, total PKR, and phospho-PKR in A549 and NCI-H1299 (**Figure 4A**), as well as NCI-H460 and NCI-H1437 cells (**Figure S4A**). The combination of *XRN1* KO with interferon-β treatment produced the greatest increase in PKR phosphorylation that was associated with a mobility shift of phospho-PKR to higher molecular weight (**Figure 4A**).

**Figure 4.**
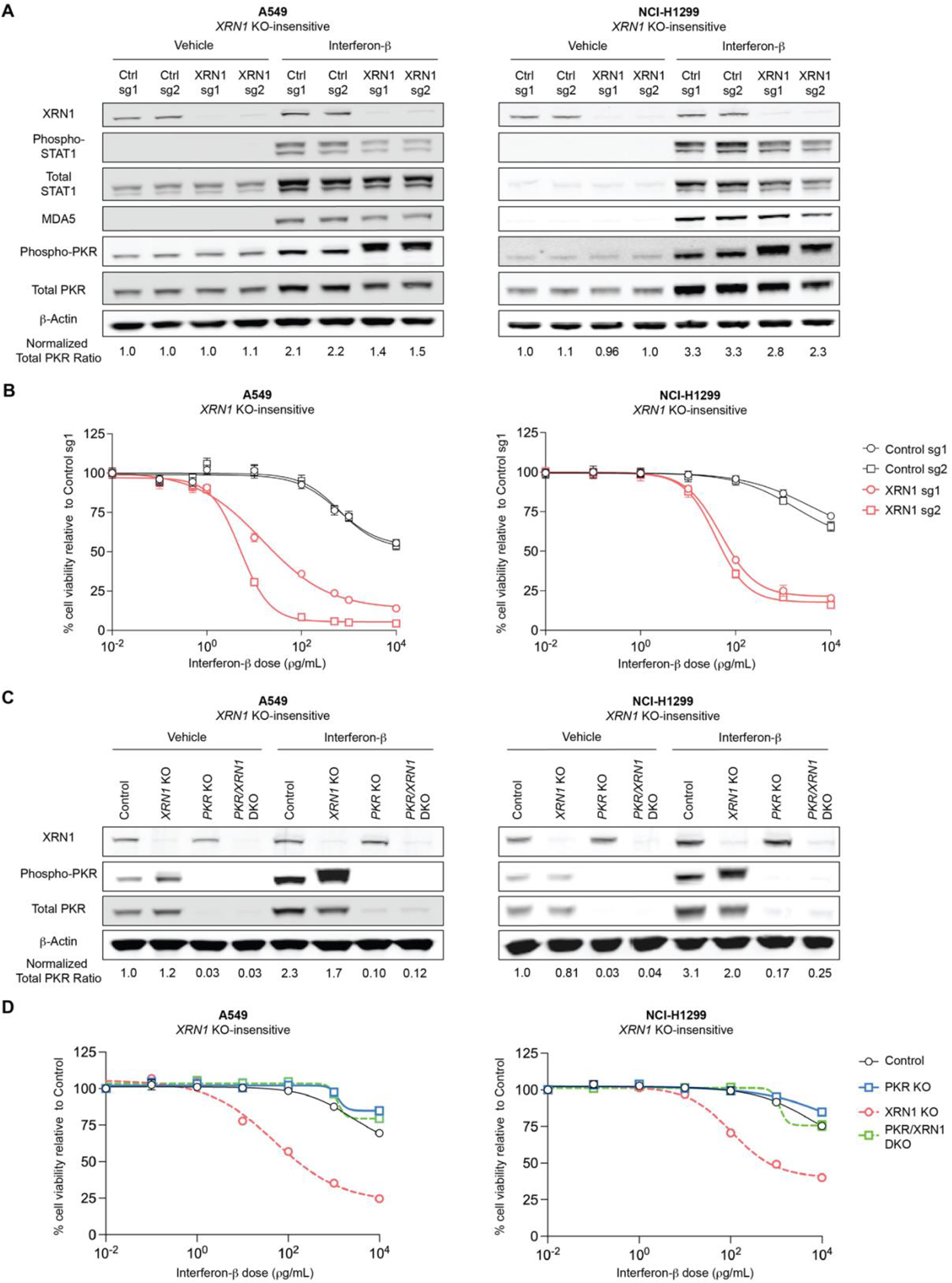
Interferon-β stimulation can increase PKR levels and sensitivity to *XRN1* deletion. (**A**) Representative immunoblots showing XRN1, phospho-STAT1, total STAT1, MDA5, phospho-PKR, total PKR, and β-actin protein levels in control or *XRN1* KO A549 (left) or NCI-H1299 (right) cells after 24 hours of treatment with vehicle control (sterile water) or interferon-β (10 ng/mL). (**B**) Cell viability was assessed by ATP bioluminescence in control or *XRN1* KO A549 (left) or NCI-H1299 (right) cells 5 days after treatment with vehicle control (sterile water) or the indicated concentrations of interferon-β. (**C**) Representative immunoblots showing XRN1, phospho-PKR, total PKR, and β-actin protein levels in control, *XRN1* single KO, *PKR* single KO, or *PKR/XRN1* double KO A549 (left) or NCI-H1299 (right) cells after 24 hours of treatment with vehicle control (sterile water) or interferon-β (10 ng/mL). (**D**) Cell viability was assessed by ATP bioluminescence in control, *XRN1* single KO, *PKR* single KO, or *PKR/XRN1* double KO A549 (left) or NCI-H1299 (right) cells 5 days after treatment with vehicle control (sterile water) or the indicated concentrations of interferon-β. Three independent biological replicates were performed for each cell line in (**A-D**).

Interferon-β stimulation of A549 and NCI-H1299 cells led to an increase in cell lethality of *XRN1*-deleted isogenic cells, compared to cells expressing control sgRNA (**Figure 4B**). This interferon-β-induced cell lethality requires PKR, as co-deletion of *PKR* and *XRN1* (**Figure 4C**) rescued cell viability at multiple doses of interferon-β (**Figure 4D**). Although interferon-β treatment activated interferon signaling in the *XRN1* KO-insensitive cell lines NCI-H460 and NCI-H1437, *XRN1* deletion did not alter either PKR phosphorylation or cell viability substantially in these cell lines (**Figures S4A** and **S4B**). These latter data suggest that additional cell line-specific factors can modulate the response to interferon signaling and *XRN1* deletion. Together, these data show that interferon signaling can induce sensitivity to *XRN1* deletion in a PKR-dependent manner and provide further support for a mechanistic connection between high ISG expression and *XRN1* genetic dependency in cancer cell lines.

### Endogenous sense/anti-sense transcripts accumulate after *XRN1* deletion

As we have shown above, the exoribonuclease activity of XRN1 is required for the survival of cancer cell lines with high ISG expression. Interferon signaling, which regulates the abundance of the key dsRNA sensor PKR, is a critical modulator of *XRN1* KO-sensitivity. Synthesizing these observations, we hypothesized that *XRN1* deletion may cause an increase in endogenous dsRNA ligands for PKR.

Thus, we examined whether *XRN1* deletion affected classes of transcripts previously reported as PKR ligands, specifically mitochondrial RNA^20^ and endogenous retroelement RNA^30^. First, we did not detect any significant increases in mitochondrial RNA transcripts after *XRN1* deletion (**Table S1**). Next, we found that *XRN1* deletion led to increased transcript levels of multiple human endogenous retrovirus 9 (HERV9) subfamilies (**Figure S5A** and **S5B; Table S3**). However, these increased HERV9 transcripts were entirely dependent on PKR activity (**Figure S5B** and **S5C; Table S3**), suggesting that HERV9 transcripts likely do not initiate PKR activation in XRN1-deficient cells. These data provide evidence that neither mitochondrial RNA nor endogenous retroelements are the culprit dsRNA ligands that stimulate PKR activation in *XRN1* KO-sensitive cancer cell lines.

An alternative mechanism for PKR activation is suggested by prior studies which showed that XRN1 regulates the formation and accumulation of complementary sense/anti-sense RNA transcripts in *S. cerevisiae*^31–33^ and during viral infection^25, 26^. To identify candidate dsRNA ligands for PKR, we performed strand-specific RNA-sequencing of *XRN1*-deleted and control isogenic cell lines to determine whether XRN1 modulates endogenous sense/anti-sense RNA transcript levels in human cancer cell lines. We developed a computational pipeline to identify regions of the transcriptome that may contain complementary sense/anti-sense RNA pairs and to quantify the abundance of these RNA pairs (**Figure 5A**). Transcriptome-wide analysis showed that targeting the *XRN1* KO-sensitive HCC366 cell line with *XRN1* sgRNA caused a significant increase in the fraction of complementary sense/anti-sense RNA pairs among all transcripts, compared to control sgRNA (**Figure 5B, Table S4**). Next, we investigated whether PKR is required for the accumulation of these complementary RNA pairs. We found that co-deletion of both *PKR* and *XRN1* caused a smaller increase in complementary sense/anti-sense RNA pairs compared to *XRN1* deletion alone (**Figure 5C; Table S4**). This result suggests that there could be two classes of XRN1-regulated complementary RNA pairs: one class that is regulated directly by XRN1 (“directly XRN1-regulated,” shown in purple) and a second class that is regulated indirectly by XRN1 via PKR activation (“PKR-regulated,” shown in green) (**Figure 5D, Table S4**). The HERV9 transcripts, described above, are an example of PKR-regulated transcripts.

**Figure 5.**
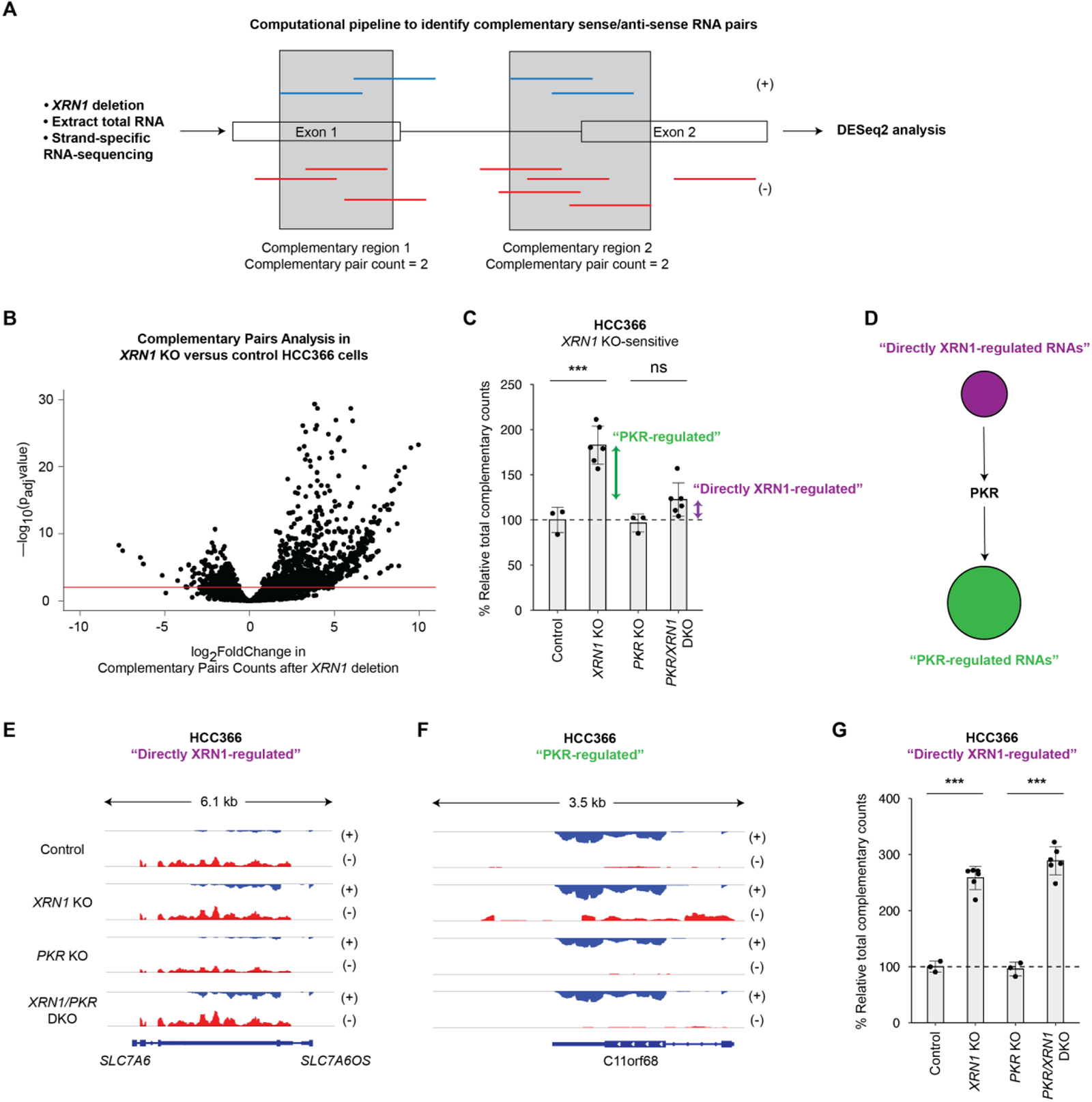
Endogenous sense/anti-sense transcripts accumulate after *XRN1* deletion. (**A**) Schematic of computational pipeline to identify complementary sense/anti-sense RNA pairs in strand-specific RNA sequencing data. See STAR Methods for additional details. (**B**) Volcano plot of differentially expressed complementary sense/anti-sense RNA pairs in *XRN1* KO compared to control HCC366 cells. RNA extraction was performed 7 days after CRISPR-Cas9 targeting of control loci or *XRN1*. The horizontal red line corresponds to a p_adj_ of 0.1. (**C**) Quantitation of relative levels of complementary sense/anti-sense RNA pairs in control, *XRN1* single KO, *PKR* single KO, and *PKR/XRN1* double KO HCC366 cells. **GREEN** arrows show the relative quantity of “PKR-regulated” complementary pairs and **PURPLE** arrows show the relative quantity of “directly XRN1-regulated” complementary pairs. RNA extraction was performed 7 days after CRISPR-Cas9 targeting of a control locus, *XRN1*, *PKR*, or both *XRN1* and *PKR*. (**D**) Schematic depicting the relationship of the dsRNA sensor PKR to “directly XRN1-regulated” and “PKR-regulated” complementary sense/anti-sense RNA pairs identified in (**C**). (**E, F**) Representative pileups from strand-specific RNA-sequencing of control, *XRN1* single KO, *PKR* single KO, and *PKR/XRN1* double KO HCC366 cells showing examples of “directly XRN1-regulated” and “PKR-regulated” complementary sense/anti-sense RNA pairs. (**G**) Quantitation of relative levels of “directly XRN1-regulated” complementary sense/anti-sense RNA pairs in control, *XRN1* single KO, *PKR* single KO, and *PKR/XRN1* double KO HCC366 cells. Relative complementary sense/anti-sense values were normalized to the control HCC366 sample. Error bars represent standard deviation. *** *p* < 0.001 and ns = not significant as calculated by two-sided Student’s *t*-test.

Among complementary sense/anti-sense transcripts classified as directly XRN1-regulated, the *SLC7A6/SLC7A6OS* (**Figure 5E**) and *HYI/SZT* (**Figure S5D**) loci represent two of the top ten most abundant RNA pairs that are significantly increased after *XRN1* deletion. Likewise, among complementary sense/anti-sense transcripts classified as PKR-regulated, the *C11orf68* (**Figure 5F**) and *CFLAR* (**Figure S5E**) loci represent two of the top ten most abundant RNA pairs that are significantly increased after *XRN1* deletion. We reasoned that the directly XRN1-regulated complementary RNA pairs may function as the RNA ligands that initially activate PKR in XRN1-deficient cancer cells. Notably, we found significantly increased levels of directly XRN1-regulated sense/antisense RNA pairs in XRN1-depleted HCC366 cells as compared to HCC366 cells expressing control sgRNA (**Figure 5G**). Taken together, these data suggest that XRN1 regulates the accumulation of endogenous complementary sense/anti-sense RNA pairs in human cancer cells, likely through both direct RNA degradation and modulation of PKR activation. Furthermore, these endogenous RNA pairs have the potential to form dsRNA through base pair complementarity and may represent candidate ligands for PKR in cancer cells.

## Discussion

Growing evidence suggests that targeting innate immune pathways in cancer cells, including regulators of dsRNA metabolism, may represent a promising therapeutic strategy^14^. Previous work from our group and others has identified the RNA editing enzyme, ADAR1, as a unique genetic dependency in a subset of cancer cells with interferon response pathway activation^7, 8, 12^. These studies have also implicated a critical role for the dsRNA sensor PKR in modulating cancer cell survival after ADAR1 inactivation.

Here, we show that the 5’ to 3’ exoribonuclease XRN1 is a selective genetic dependency in human patient-derived cancer cells, with *XRN1* dependency correlated to activation of the interferon response pathway. We demonstrate further that human cancer cell lines with an activated interferon cell state tend to have higher levels of the dsRNA sensor PKR and that PKR is required for *XRN1* genetic dependency.

We propose that XRN1 activity allows cancer cells to tolerate activation of the interferon response pathway as described in the following model (**Figure S5F**). One role for XRN1 is to degrade anti-sense RNA transcripts, thereby preventing formation of complementary sense/anti-sense RNA pairs and limiting activation of the PKR dsRNA sensing pathway. This proposed mechanism is concordant with observations in yeast, where XRN1-sensitive unstable transcripts can form dsRNA structure^32, 33^. Depletion of XRN1 in interferon-activated cancer cells causes accumulation of complementary sense/anti-sense RNA pairs, which may function as dsRNA ligands for PKR that trigger PKR activation and cancer cell lethality. Disruption of the activated interferon cell state with the JAK1/2 inhibitor ruxolitinib can decrease cellular PKR levels and rescue cell lethality induced by XRN1 depletion. Conversely, stimulation of cancer cell lines with lower expression of ISGs with exogenous interferon-β can increase cellular PKR levels and induce sensitivity to XRN1 depletion. Overall, our data suggest that the abundance of both dsRNA ligand and PKR levels may contribute to *XRN1* KO-sensitivity in cancer cells with interferon pathway activation.

We showed that XRN1 regulates the accumulation of endogenous complementary sense/anti-sense RNA pairs, which may represent candidate PKR ligands. Studies using crosslinking and immunoprecipitation followed RNA-sequencing (CLIP-seq) analysis of PKR in XRN1-sufficient cells have identified bi-directionally transcribed mitochondrial RNAs and inverted *Alu* retroelements as bona fide PKR ligands^20^. Whether PKR is activated by specific complementary RNA pairs or a general accumulation of sequence-agnostic complementary RNA pairs in *XRN1* KO-sensitive cancer cells remains an open question. Answering this complex question is likely to require the simultaneous targeting of multiple genomic loci that encode complementary RNA pairs, requiring the development of novel experimental methods.

Viewing the regulation of dsRNA more broadly, a recent study showed that sense/anti-sense RNAs may be potential substrates for human ADAR1^34^, which is also a negative regulator of PKR. In addition, analysis of genome-wide association studies showed that a subset of these ADAR1-edited complementary RNA pairs were transcribed at genomic loci implicated in the pathogenesis of a range of autoimmune diseases^34^. Thus, future studies may reveal additional roles for complementary sense/anti-sense RNA pairs in cancer and autoimmune disease.

A fraction of the endogenous complementary sense/anti-sense RNA pairs we identified in XRN1-deficient cells were PKR-regulated. Prior studies have shown that DNA virus infection can trigger host RNA polymerases to transcribe immunogenic dsRNA from the viral genome^35, 36^, thereby amplifying the antiviral innate immune response. Similarly, our data suggest that *XRN1* deletion can activate PKR to cause the accumulation of additional PKR-regulated complementary sense/anti-sense RNA pairs. These PKR-regulated complementary RNA pairs may serve as a positive feedback mechanism to amplify PKR activation. Whether PKR signaling can generate “endogenous adjuvant” signals for the innate immune system merits additional investigation.

Our work suggests that interferon pathway activation can induce *XRN1* KO-sensitivity, at least in part, by controlling the abundance of the interferon-inducible protein PKR. Of note, several studies have demonstrated that a subset of human cancer cell lines and patient tumors of diverse lineages exhibit cancer cell-autonomous interferon pathway activation^8, 37, 38^. From a therapeutic perspective, our study suggests that interferon pathway activation in cancer cells may create cell state-specific genetic vulnerabilities whereby targeting molecular regulators of dsRNA sensors, such as PKR, can stimulate a viral mimicry response to promote cancer cell lethality. Aside from genetic vulnerabilities, one study showed that the activated interferon cell state enhances the response to anthracycline-based chemotherapy in cancer cell lines and patients with breast cancer^37^. Identification of additional genetic or pharmacologic vulnerabilities associated with the activated interferon cell state may improve therapeutic targeting of this subset of human cancers.

In addition to nominating XRN1 as a potential therapeutic target to induce direct cancer cell killing, our data may have implications for overcoming resistance to cancer immunotherapy. Multiple studies have demonstrated that resistance to immune checkpoint inhibition is associated with activation of an ISG expression program in cancer cells^39–41^. While an elevated ISG expression program may confer an immunotherapy-resistant cell state, activated interferon signaling may also increase the abundance of PKR and create a cell state-specific vulnerability to XRN1 inhibition. A recent study showed that XRN1 depletion in mouse tumor models can potentiate the effect of cancer immunotherapy through a MAVS-dependent mechanism^42^. While the mouse cancer cells tested in that study do not require XRN1 for survival, future work may clarify the precise role of PKR in the response of XRN1-depleted cells to cancer immunotherapy.

Taken together, our study provides mechanistic insight into how XRN1 regulates the activity of the dsRNA sensor PKR in cancer cells. Moreover, our data nominate XRN1 as a potential therapeutic target in cancers with high expression of ISGs. Future studies will be required to determine the optimal approach to leverage XRN1 biology for cancer therapeutics.

## Acknowledgements

The authors thank the members of the Meyerson laboratory for helpful discussions, Elisa Izaurralde for generously sharing the *XRN1* ORF (Addgene), and Elizabeth Henske and Damir Khabibulin for generous access to their confocal microscope. This research was supported by NIH grants, T32 CA009172 (T.Z.), K08 CA252169 (T.Z.), and R35 CA197568 (M.M), the American Society of Clinical Oncology Endowed Young Investigator Award in memory of John R. Durant, M.D. (T.Z.), and the American Cancer Society Research Professorship (M.M).

## Author Contributions

T.Z. and M.M. conceived of the study. T.Z. designed and performed most of the experiments and analyzed most of the data. M.Z. conceived and wrote the source code for the complementary sense/anti-sense RNA pairs computational pipeline. P.Z., A.R.F., and H.Y.W. performed experiments and analyzed data. A.G, P.Z., Z.Z., and A.D.C. performed computational analysis. T.Z., M.Z., A.G., and M.M. composed the manuscript. All authors read and edited the manuscript.

## Declaration of Interests

M.Z. is now an employee of Bayer Pharmaceuticals in Cambridge, MA. M.M. receives research funding from Bayer Pharmaceuticals and Janssen Pharmaceuticals; has a consulting role and equity with Delve Bio, Interline, and Isabl; and receives patent royalties on intellectual property from The Broad Institute of Harvard and MIT and Dana-Farber Cancer Institute licensed to Bayer and LabCorp, respectively. The remaining authors declare no competing interests.

---

**Supplemental Figure 1.**
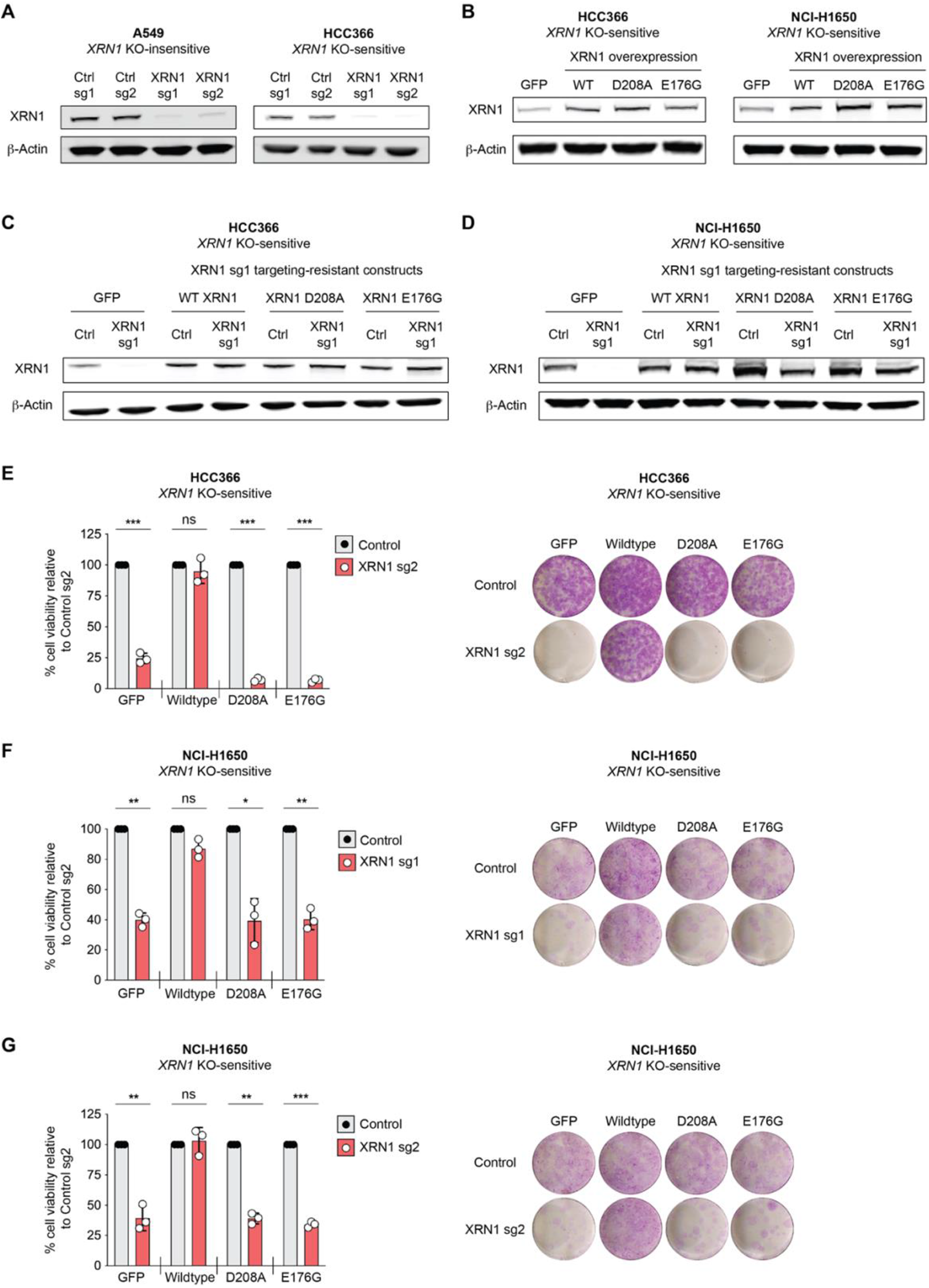
The enzymatic activity of XRN1 is required for survival of *XRN1* KO-sensitive cancer cell lines. (**A**) Representative immunoblots confirming specific depletion of XRN1 after CRISPR-Cas9 targeting of control loci or *XRN1*. β-actin was used as a loading control. (**B**) Immunoblots showing XRN1 protein levels after expression of *GFP* or overexpression of WT or catalytically inactive mutant versions of *XRN1* that are resistant to XRN1 sg1/2 targeting by CRISPR-Cas9. β-actin was used as a loading control. (**C, D**) Immunoblots showing XRN1 protein levels after CRISPR-Cas9 targeting of a control locus or *XRN1* in HCC366 (**C**) and NCI-H1650 (**D**) cells expressing *GFP* or overexpressing WT or catalytically inactive mutant versions of *XRN1* that are resistant to XRN1 sg1 targeting by CRISPR-Cas9. β-actin was used as a loading control. (**E**) Cell viability was assessed by ATP bioluminescence (left panel) or crystal violet staining (right panel) at 12 or 17 days, respectively, after targeting of a control locus or endogenous *XRN1*, by XRN1 sg2, with CRISPR-Cas9 in **HCC366** cells expressing *GFP* control or overexpressing WT *XRN1* resistant to *XRN1* sg2 targeting or catalytically inactive mutants of *XRN1* (D208A and E176G) resistant to *XRN1* sg2 targeting. (**F, G**) Cell viability was assessed by ATP bioluminescence (left panels) or crystal violet staining (right panels) at 12 or 21 days, respectively, after targeting of a control locus or endogenous *XRN1*, by XRN1 sg1 in (**F**) and sg2 in (**G**), with CRISPR-Cas9 in **NCI-H1650** cells expressing *GFP* control or overexpressing WT *XRN1* resistant to XRN1 sg1/2 targeting or catalytically inactive mutants of *XRN1* (D208A and E176G) resistant to *XRN1* sg1/2 targeting. ATP bioluminescence values were normalized to the control sg1 sample within each isogenic cell line. Data from three independent biological replicates are shown in (**E**-**G**). Error bars represent standard deviation. \**p* < 0.05, \*\**p* < 0.01, *** *p* < 0.001, and ns = not significant as calculated by paired Student’s *t* test in (**E-G**).

**Supplemental Figure 2.**
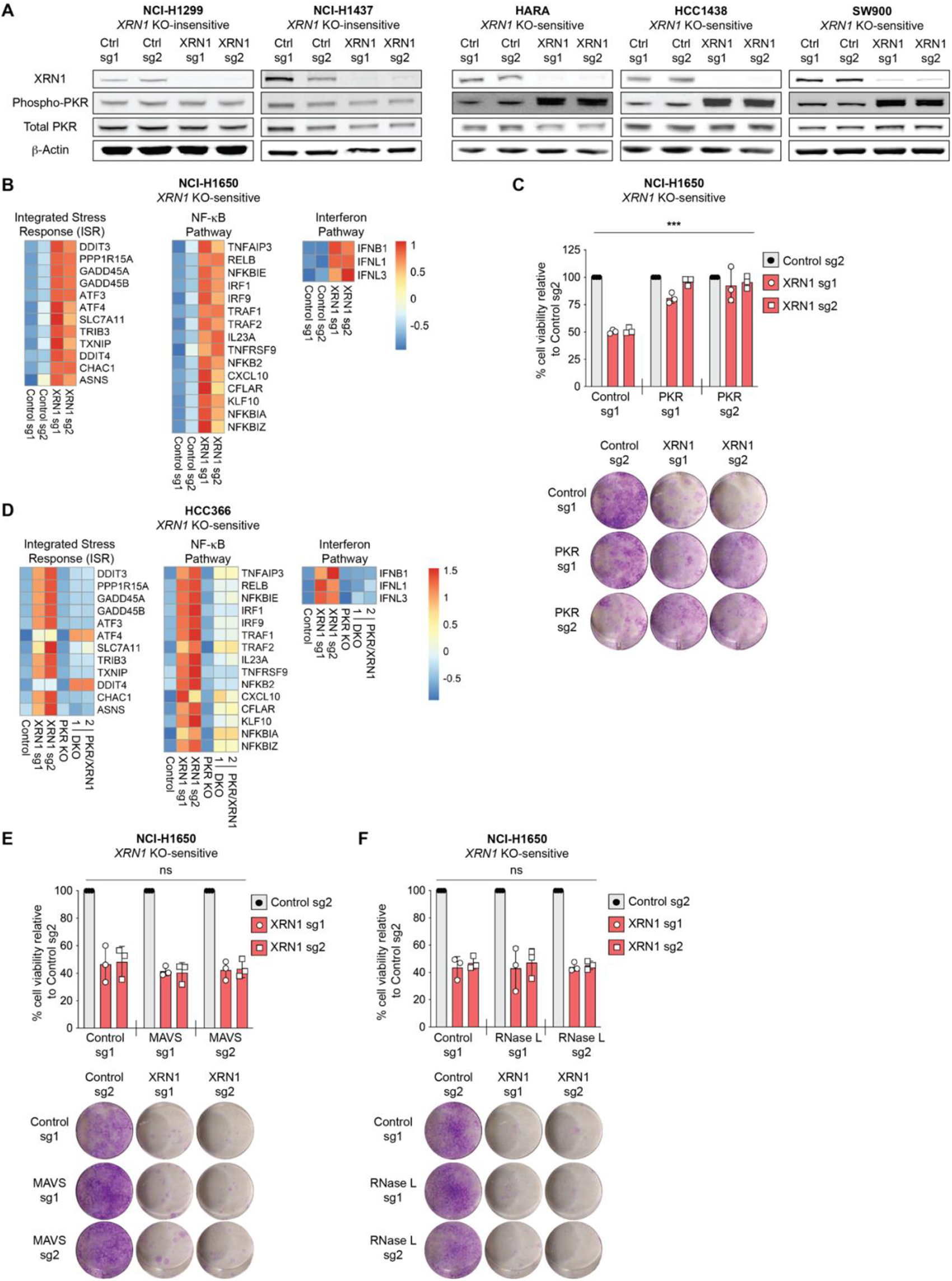
PKR signaling is required for *XRN1* gene essentiality. (**A**) Representative immunoblots showing XRN1, phospho-PKR, total PKR, and β-actin protein levels in *XRN1* KO-insensitive (NCI-H1299 and NCI-H1437) and *XRN1* KO-sensitive (HARA, HCC1438, and SW900) cells 7 days after CRISPR-Cas9 targeting of control loci or *XRN1*. At least three independent biological replicates were performed for each cell line. (**B**) Heatmaps showing relative TPM values for the indicated differentially expressed genes in the integrated stress response, NF-κB, and interferon pathways (rows) 7 days after CRISPR-Cas9 targeting of control loci or *XRN1* (columns) in **NCI-H1650** cells. **RED**: increased expression, **BLUE**: decreased expression, as shown on the right side of the diagram. Each condition includes three independent biological replicates. (**C**) (Top panel) Cell viability was assessed by ATP bioluminescence 12 days after CRISPR-Cas9 targeting of a control locus or *XRN1* in control-targeted or PKR-depleted **NCI-H1650** cells. (Bottom panel) Crystal violet staining 21 days after CRISPR-Cas9 targeting of a control locus or *XRN1* in control-targeted or PKR-depleted NCI-H1650 cells. (**D**) Heatmaps showing relative TPM values for the indicated differentially expressed genes in the integrated stress response, NF-κB, and interferon pathways (rows) 7 days after CRISPR-Cas9 targeting of a control locus, *XRN1*, *PKR*, or both *XRN1* and *PKR* (columns) in **HCC366** cells. **RED**: increased expression, **BLUE**: decreased expression, as shown on the right side of the diagram. Each condition includes at least three independent biological replicates. (**E, F**) (Top panel) Cell viability was assessed by ATP bioluminescence 12 days after CRISPR-Cas9 targeting of a control locus or *XRN1* in control-targeted or MAVS-depleted (**E**) or control-targeted or RNase L-depleted (**F**) **NCI-H1650** cells. (Bottom panel) Crystal violet staining 21 days after CRISPR-Cas9 targeting of a control locus or *XRN1* in control-targeted or MAVS-depleted (E) or control-targeted or RNase L-depleted (**F**) **NCI-H1650** cells. In (**C, E, F**), ATP bioluminescence values were normalized to the control sg1 sample within each cell line. Data from three independent biological replicates are shown. Error bars represent standard deviation. *** *p* < 0.001 and ns = not significant, as calculated by repeated measures two-way ANOVA. Crystal violet images are representative of three independent biological experiments.

**Supplemental Figure 3.**
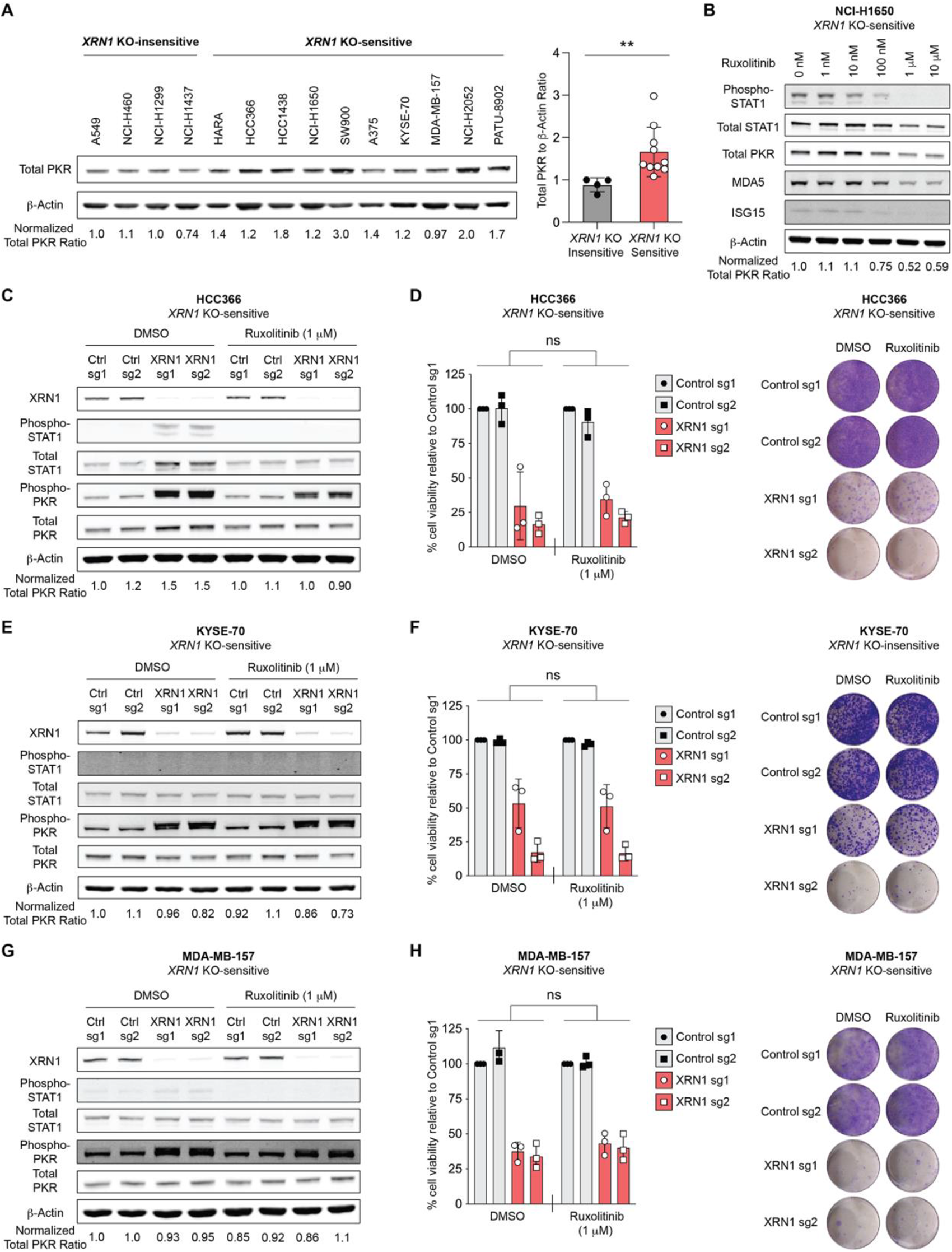
Inhibition of JAK1/2 signaling does not affect PKR levels or *XRN1* KO-sensitivity in XRN1 KO-sensitive cell lines with lower baseline phospho-STAT1 levels. (**A**) (Left panel) Representative immunoblot showing total PKR protein levels in parental *XRN1* KO-insensitive and KO-sensitive cancer cell lines. β-actin was used as a loading control. (Right panel) Bar graph showing total PKR protein levels normalized to β-actin from parental *XRN1* KO-insensitive and KO-sensitive cancer cell lines shown in (**A**). Each dot represents the averaged normalized total PKR to β-actin ratio of an individual cell line across three independent biological experiments. ** < 0.01 as calculated by Welch’s *t*-test. (**B**) Immunoblot showing phospho-STAT1, total STAT1, total PKR, MDA5, ISG15 and β-actin protein levels in *XRN1* KO-sensitive NCI-H1650 cells treated with either DMSO control or varying concentrations of ruxolitinib for 24 hours. (**C**, **E**, **G**) Representative immunoblots showing XRN1, phospho-STAT1, total STAT1, phospho-PKR, total PKR, and β-actin protein levels in control and *XRN1*-deleted HCC366 (**C**), KYSE-70 (**E**), and MDA-MB-157 (**G**) cells treated with either DMSO control or ruxolitinib (1μM). Three independent biological replicates were performed for each cell line. (**D**, **F**, **H**) Cell viability was assessed by either ATP bioluminescence (left panels) or crystal violet staining (right panels) after CRISPR-Cas9 targeting of control loci or *XRN1* in HCC366 (**D**), KYSE-70 (**F**), and MDA-MB-157 (**H**) cells treated with DMSO control or ruxolitinib (1μM). ATP bioluminescence values were normalized to the control sg1 sample within each cell line. Data from three independent biological replicates are shown. Error bars represent standard deviation. \**p* < 0.05 and *** *p* < 0.001, as calculated by repeated measures two-way ANOVA. Crystal violet images are representative of three independent biological experiments.

**Supplemental Figure 4.**
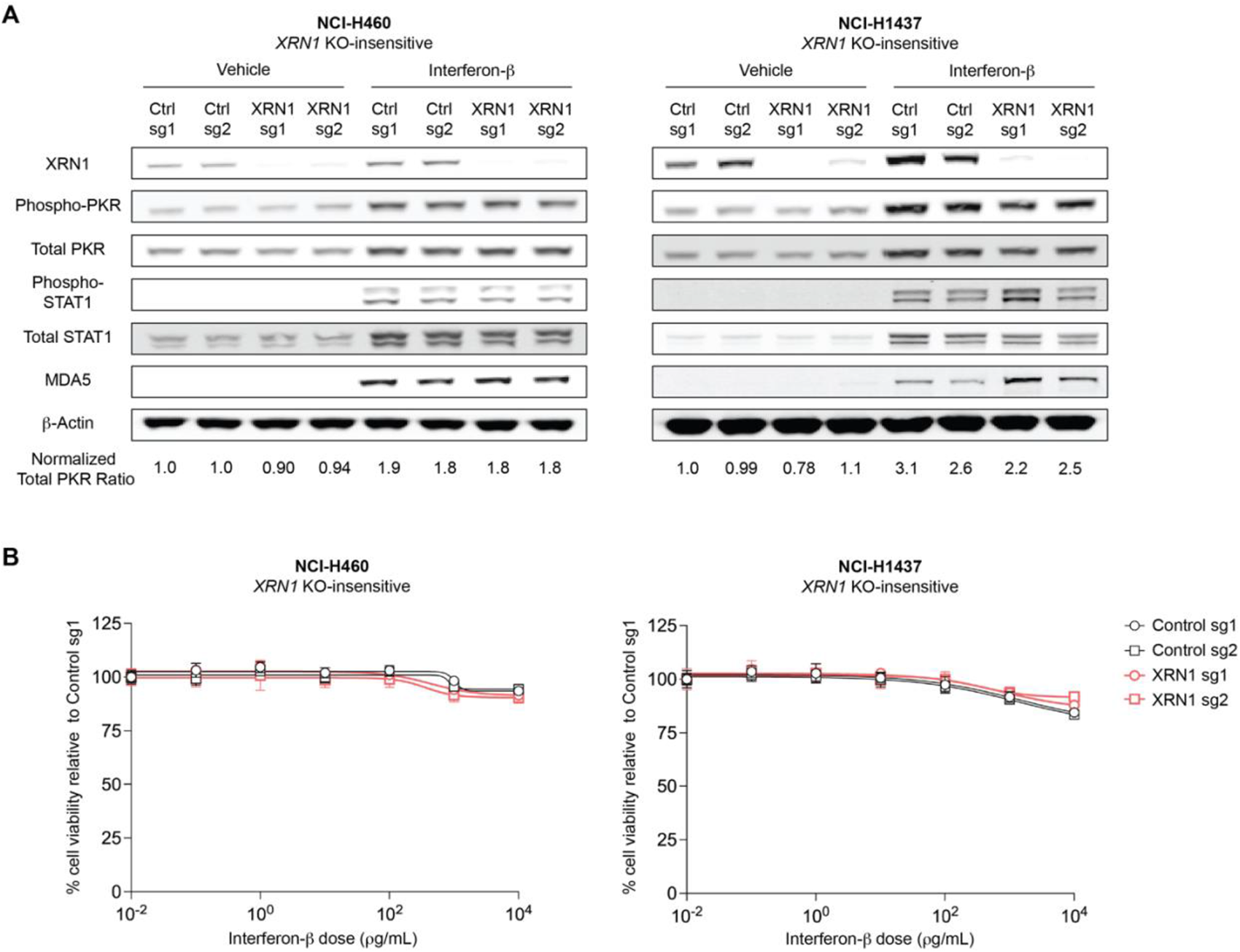
Interferon-β stimulation can increase PKR levels, but does not affect sensitivity to *XRN1* deletion in a subset of *XRN1* KO-insensitive cell lines. (**A**) Representative immunoblots of XRN1, phospho-STAT1, total STAT1, MDA5, phospho-PKR, total PKR, and β-actin protein levels in control or *XRN1* KO NCI-H460 (left) or NCI-H1437 (right) cells after 24 hours of treatment with vehicle control (sterile water) or interferon-β (10 ng/mL). (**B**) Cell viability was assessed by ATP bioluminescence in control or *XRN1* KO NCI-H460 (left) or NCI-H1437 (right) cells 5 days after treatment with vehicle control (sterile water) or the indicated concentrations of interferon-β. Three independent biological replicates were performed for each cell line.

**Supplemental Figure 5.**
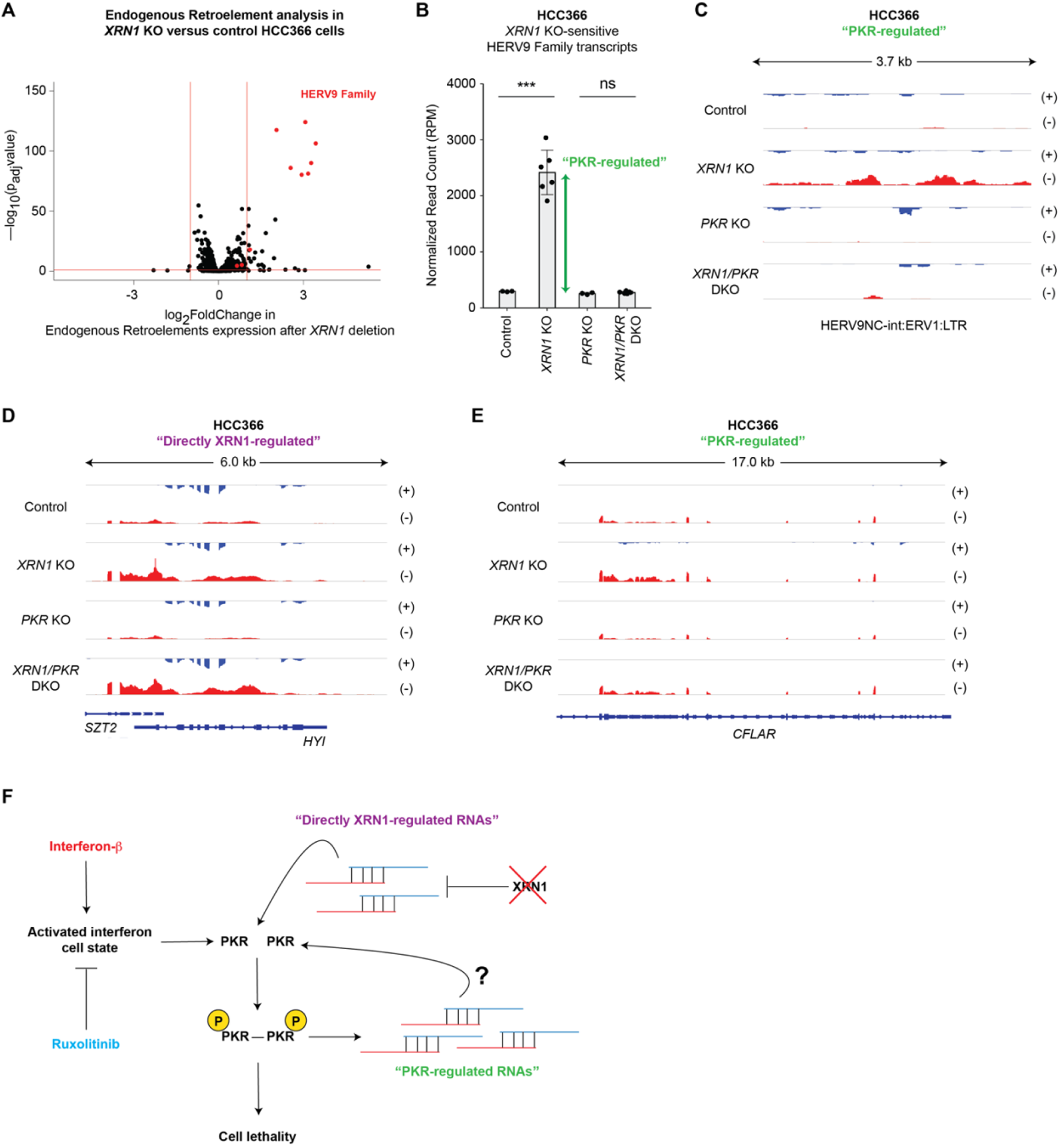
Endogenous sense/anti-sense transcripts accumulate after *XRN1* deletion. (**A**) Volcano plot of differentially expressed endogenous retroelement subfamilies in *XRN1* KO compared to control HCC366 cells. RNA extraction was performed 7 days after CRISPR-Cas9 targeting of control loci or *XRN1*. The SQuIRE computational pipeline was used to identify endogenous retroelements. Each dot corresponds to a retroelement subfamily. The horizontal red line corresponds to a p_adj_ of 0.1. The vertical red lines correspond to a two-fold decrease (left) and a two-fold increase (right) in expression levels. (**B**) Quantitation of HERV9 family RNA transcripts in control, *XRN1* single KO, *PKR* single KO, and *PKR/XRN1* double KO HCC366 cells. **GREEN** arrow represents the approximate quantity of PKR-regulated HERV9 family transcripts. RNA extraction was performed 7 days after CRISPR-Cas9 targeting of a control locus, *XRN1*, *PKR*, or both *XRN1* and *PKR*. Normalized read counts of all annotated HERV9 family transcripts are shown. Error bars represent standard deviation. *** *p* < 0.001 and ns = not significant as calculated by Student’s *t*-test. (**C)** Representative pileups from strand-specific RNA-sequencing of control, *XRN1* single KO, *PKR* single KO, and *PKR/XRN1* double KO HCC366 cells showing an example of a PKR-regulated HERV9 family transcript. (**D, E**) Representative pileups from strand-specific RNA-sequencing of control, *XRN1* single KO, *PKR* single KO, and *PKR/XRN1* double KO HCC366 cells showing additional examples of directly XRN1-regulated and PKR-regulated complementary sense/anti-sense RNA pairs. (**F**) Schematic model of the molecular factors that contribute to *XRN1* genetic dependency.

## Supplemental Tables

**Supplemental Table 1. Differentially expressed genes between control and *XRN1* KO HCC366 and NCI-H1650 cells.** A list of differentially expressed genes between *XRN1* KO versus control HCC366 (first tab) or NCI-H1650 (second tab) cells. Column A displays the Ensembl gene ID, column B displays the gene type, and column C displays the gene name. The associated log2-fold change values (column D), *p* values (column E), and *p*_adj_ values (column F) are provided for each differentially expressed gene.

**Supplemental Table 2. Differentially expressed genes in control, *XRN1* single KO, *PKR* single KO, and *XRN1/PKR* double KO HCC366 cells.** A list of differentially expressed genes between *XRN1* single KO versus control (first tab), *PKR* single KO versus control (second tab), and *XRN1*/*PKR* double KO versus control (third tab) HCC366 cells. Column A displays the Ensembl gene ID, column B displays the gene type, and column C displays the gene name. The associated log2-fold change values (column D), *p* values (column E), and *p*_adj_ values (column F) are provided for each differentially expressed gene.

**Supplemental Table 3. Differentially expressed endogenous retroelement subfamilies between control, *XRN1* single KO, or *XRN1*/*PKR* double KO HCC366 cells.** A list of differentially expressed endogenous retroelement subfamilies (column A) between *XRN1* KO versus control (first tab) and *XRN1*/*PKR* double KO versus control (second tab) HCC366 cells. The associated log2-fold change values (column B), *p* values (column C), and *p*_adj_ values (column D) are provided for each differentially expressed endogenous retroelement subfamily.

**Supplemental Table 4. Differentially expressed complementary sense/anti-sense RNA pairs in control, *XRN1* single KO, or *XRN1/PKR* double KO HCC366 cells.** A list of differentially expressed complementary sense/anti-sense RNA pairs between *XRN1* KO versus control (first tab) and *XRN1*/*PKR* double KO versus control (second tab) HCC366 cells. Each genomic region containing a differentially expressed complementary RNA pair is defined by its chromosomal location: chromosome number in column A, start position in column B, and end position in column C. The associated log2-fold change values (column D), *p* values (column E), *p*_adj_ values (column F), nearest gene(s) (column G), and transcript type (column H) are provided for each complementary RNA pair. Column I in the first tab indicates whether the complementary RNA pairs are categorized as directly XRN1-regulated or PKR-regulated transcripts.

**Supplemental Table 5.**
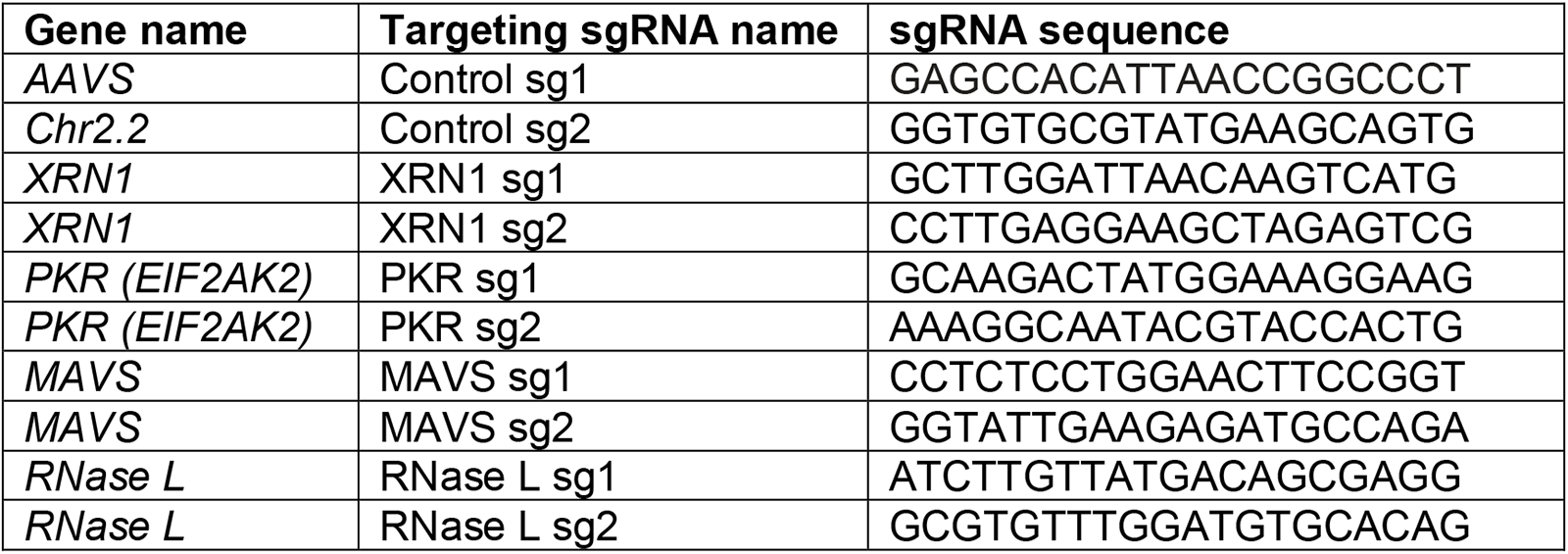
sgRNA sequences utilized for CRISPR-Cas9-mediated gene deletion. The targeted gene (column A), name of the sgRNA used in this study (column B), and sgRNA sequence (column C) are provided.

## Materials and Methods

### Resource availability

#### Lead contact

Further information and requests for reagents may be directed to and will be fulfilled by the corresponding author, Matthew Meyerson.

#### Materials availability

All plasmids generated in this study will be deposited in Addgene upon publication. Any additional materials will be available upon publication and request.

#### Data and code availability

The raw and processed bulk RNA-seq data presented in this manuscript will be deposited at the NCBI Gene Expression Omnibus (GEO) repository (deposite pending) and will be publicly available as of the date of publication. Access numbers are listed in the key resources table. The source data used to generate **Figures 1A-D** and **3A** were obtained from publicly available datasets from the Cancer Dependency Map Portal [http://depmap.org/portal/]. Microscopy data reported in this paper will be shared by the lead contact upon request.

All original code in this manuscript has been deposited at GitHub and is publicly available. The DOI to access the original code is listed in the key resources table.

Any additional information required to re-analyze the data reported in this paper will be available from the lead contact upon request.

### Experimental model and subject details

#### Cell lines

A549 (Male), NCI-H460 (Male), NCI-H1299 (Male), NCI-H1437 (Male), HARA (Male), HCC366 (Female), HCC1438 (Male), NCI-H1650 (Male), KYSE-70 (Male), MDA-MB-157 (Female), and NCI-H2052 (Male) were cultured in RPMI supplemented with 10% fetal bovine serum. A375 (Female), PATU-8902 (Female), and HEK 293T (Female) cells were cultured in DMEM supplemented with 10% fetal bovine serum. All cells were cultured at 37 degrees Celsius with 5.0% carbon dioxide.

All cancer cell lines were obtained from the Broad Institute Cancer Cell Line Encyclopedia^44^ (CCLE) or the American Type Culture Collection (ATCC). Both CCLE and ATCC perform cell line authentication and mycoplasma testing routinely. All cell lines were tested for mycoplasma prior to use and regularly thereafter.

### Method details

#### CRISPR-Cas9 gene knockout

Single guide RNA sequences were designed using the sgRNA Designer tool (The Broad Institute Genomics Perturbation Platform) (https://portals.broadinstitute.org/gpp/public/analysis-tools/sgrna-design). sgRNA sequences are displayed in **Table S5**. sgRNAs were cloned into the Cas9 expressing lentiviral vector lentiCRISPRv2 or a modified lentiCRISPRv2 construct that expresses two different sgRNAs under the control of separate human and mouse U6 promoters. Individual lentiCRISPRv2 vectors were introduced along with packaging vectors into human embryonic kidney (HEK) 293T cells via calcium phosphate transfection according to the manufacturer’s protocol (Clontech). Lentivirus was harvested at 48 and 72 hours after transfection in RPMI media supplemented with 10% FBS and filtered with 45 μm filters prior to transduction of cancer cell lines using centrifugation at 1000 *g* for 2 hours in the presence of 8 μg/mL of polybrene (Santa Cruz Biotechnology). Transduced cell lines were selected in puromycin and/or blasticidin (Thermo Fisher Scientific) for at least 5 days prior to use in assays. Protein lysates were collected from the transduced cells and protein levels of the targeted gene(s) were assessed by immunoblotting to validate gene KO.

#### Cell viability assays

Cell counting was performed using a Coulter Particle Counter (Beckman-Coulter). For ATP bioluminescence experiments, cells were plated at a density of 1,500 (HARA, HCC1438, NCI-H2052, PATU-8902) or 3,000 (A375, A549, KYSE-70, MDA-MB-157, NCI-H460, NCI-H1299, NCI-H1437, HCC366, NCI-H1650) cells per well in 96 well assay plates (Corning). ATP bioluminescence was assessed at 12 days after gene KO with the CellTiter-Glo Luminescent Cell Viability Assay (Promega). For crystal violet staining, cells were plated at a density of 10,000 or 20,000 cells per well in 12 well tissue culture plates. Once the control cells grew to near confluency, each well was washed twice with ice cold PBS, fixed on ice with ice cold methanol for 10 minutes, stained with 0.5% crystal violet solution (made in 25% methanol) for 10 minutes at room temperature, and washed at least four times with water. All cell viability assays were performed in at least triplicate.

#### XRN1 mutagenesis and overexpression

An *XRN1* open-reading frame (ORF) clone deposited by Elisa Izaurralde was purchased from Addgene (#66596). The entire ORF was sequenced to confirm fidelity to the NCBI Reference Sequence NM_019001.5. An entry clone for *XRN1* was obtained through PCR-amplification of the *XRN1* ORF and was sub-cloned into a Gateway donor vector. Overlap PCR was performed to introduce silent mutations separately into the protospacer adjacent motif (PAM) sequences targeted by XRN1 sgRNA 1 and sgRNA 2, thereby rendering the constructs resistant to CRISPR-Cas9 editing by these sgRNAs. Subsequently, the D208A and E176G mutations were engineered into each XRN1 sg1 and sg2-resistant *XRN1* construct separately using overlap PCR. The resulting XRN1 sg1 and sg2-resistant *XRN1* constructs and a *GFP* control construct were sub-cloned into the pLEX307 lentiviral expression vector (Addgene) under the control of an EF-1α promoter. Each expression vector was then transfected into HEK293T cells to generate lentivirus. Lentiviral transduction of target cell lines was performed as described above.

#### Immunoblotting

Cells were lysed in RIPA lysis buffer (Thermo Fisher Scientific) supplemented with 1× protease and phosphatase inhibitor cocktails (Roche). Protein concentrations were obtained using the BCA Protein Assay Kit (Pierce) and 6X Laemmli SDS sample buffer (Thermo Fisher Scientific) was added to protein extracts. Protein extracts were normalized between all samples within an experiment and boiled above 95 degrees Celsius for 10 minutes. Proteins were resolved on 4-12% Bis-Tris gradient gels, transferred to nitrocellulose membranes, and immunoblotting with primary and secondary antibodies was performed according to standard procedures. All primary antibodies were used at a dilution of 1:1000 except β-actin which was used at a dilution of 1:10,000. Secondary antibodies from LI-COR Biosciences were used at a dilution of 1:10,000. Selected immunoblots were stripped with Restore PLUS Western Blot Stripping buffer (Thermo Fisher Scientific, #21059) prior to repeat immunoblotting. Immunoblots were imaged using the LI-COR digital imaging system. Quantitation of band intensities was performed with ImageJ. All immunoblots were cropped to optimize clarity and presentation.

#### Fluorescence microscopy

Control and *XRN1* KO HCC366 cells were seeded on coverslips to 60-80% confluency. Cells were fixed with 4% paraformaldehyde for 10 minutes at room temperature, washed with PBS, and permeabilized with 0.2% Triton-X for 10 minutes at room temperature. Fixed and permeabilized cells were incubated with phosphate buffered saline with Tween (PBS-T) containing 1% bovine serum albumin for 30 minutes at room temperature and washed with PBS-T. Samples were stained using primary antibody for 1 hour at room temperature, washed twice with PBS-T, stained with secondary antibody for 1 hour at room temperature, and then washed twice with PBS-T. Nuclei were stained with Hoechst 33342 for 5 minutes at room temperature. Coverslips were mounted using Fluoromount-G and imaged with an Olympus Fluoview FV10i confocal microscope at 60X magnification with an oil-immersion lens. Microscopy images were processed with ImageJ.

#### Ruxolitinib treatment

Parental cancer cell lines were plated in 6 well plates and treated with DMSO or ruxolitinib the next day. After 24 hours of ruxolitinib treatment, protein lysates for each parental cell line were harvested as described above. Control and *XRN1* KO cancer cell lines were treated with DMSO or ruxolitinib starting at 24 hours after lentiviral transduction. Culture media containing DMSO or ruxolitinib was changed at least once every 3 days until protein lysates were harvested or cell viability measurements were obtained.

#### Interferon treatment

For interferon treatment assays, cells were plated at a density of 3,000 cells per well in a 96 well assay plate (Corning). The following day, cells were treated with increasing doses of recombinant human IFN-beta 1a (mammalian) protein (PBL Assay Science, #114151) or sterile water as the vehicle control. Cell viability was assessed 5 days after interferon-β treatment with CellTiter-Glo Luminescent Cell Viability Assay (Promega). ATP bioluminescence values of the interferon-β treated wells were normalized to those of the vehicle-treated controls. Dose curves were obtained using nonlinear regression on a standard four-parameter logistic model using GraphPad Prism 7.

#### Sample preparation and RNA-sequencing

RNA was isolated from HCC366 and NCI-H1650 cells 7 days after transduction with lentivirus co-expressing Cas9 and sgRNAs targeting control loci, *XRN1*, *PKR*, or both *XRN1* and *PKR*. RNA was isolated using the RNeasy Plus Kit (Qiagen) followed by ribosomal RNA depletion using the NEBNext rRNA Depletion Kit (E6310) or the QIAseq FastSelect -rRNA HMR Kit (Qiagen). RNA sequencing libraries were prepared using the NEBNext Ultra II Directional RNA Library Prep Kit (E7760) and sequenced on an Illumina HiSeq instrument (150-bp paired-end reads).

#### Processing of RNA-sequencing data

Quality check of RNA-sequencing FASTQ files was completed using FastQC v0.11.9 with default parameters. After manual inspection, it was determined that the first three nucleotides of all reads should be removed before mapping. Adapter trimming was completed using cutadapt v2.8; Illumina Universal Adapter sequence “AGATCGGAAGAG” was used for both ends. After trimming, reads shorter than 35 bp were discarded. Reads were mapped to the Ensembl v99 hg38 gene annotation using the STAR aligner v2.7.5b. Read pileup was extracted using STAR.

#### Differential gene expression analysis of RNA-sequencing data

RSEM^45^ was also run on aligned BAM files with parameters --estimate-rpsd --seed 12345 -- strandedness reverse to compute TPMs for each gene. Gene expression heatmaps were generated using TPM values and the R package pheatmap [https://cran.r-project.org/web/packages/pheatmap/index.html]. DESeq2^46^ was used to evaluate for differential gene expression by estimation of fold change and dispersion of data. The cutoff for differential expression between two groups was a false discovery rate (FDR) < 0.1. Fold changes for each gene were calculated by the average log2 expression value across groups, where positive values represent upregulation and negative values represent downregulation of gene expression.

#### Identification of endogenous retroelements in RNA-sequencing data

We used the SQuIRE^47^ computational pipeline to quantify endogenous retroelements at the subfamily level. SQuIRE was run with default parameters with the steps including Build, Fetch, and Map. The Count step was run twice for each sample, with the strandedness parameters set to 1 and 2 to obtain the read counts on the sense and reverse strands, respectively.

#### Transcriptome-wide identification of complementary sense/anti-sense RNA pairs

We defined bi-directionally transcribed loci as genomic regions that are covered by at least one RNA-sequencing read on both strands. In this step, pair-end reads were converted to fragments by joining the two mates and setting its orientation following the first mate. If one of the two mates was not mapped, the other mate was still included in the analysis. For simplicity, all such mapped read singletons or fragments were referred to as “reads” in this section. To be considered as a bi-directionally transcribed locus, a region must be supported by at least two reads, one mapped to the forward strand and the other to the reverse strand. We named the RNA transcripts derived from these bi-directionally transcribed loci as “complementary sense/anti-sense RNA pairs.”

Many of these regions resulted from overlapping portions of only two reads, leading to a heavily fragmented bi-directional transcriptome for any one RNA-seq sample. To enable effective quantitative analysis between different experimental conditions, we constructed union sets of these regions across multiple samples. Regions belonging to samples from the same cancer cell line were used to construct one union set by merging all overlapping regions. We then quantified the abundance of bi-directional transcription in each sample based on the number of reads mapped within the range of each merged region.

#### Quantifying the abundance of complementary sense/anti-sense RNA pairs

To represent the abundance of bi-directional transcription within a region, we defined a metric termed complementary pairs (CP). A complementary pair consists of two overlapping reads that were mapped to the forward and reverse strands, respectively. The CP value of a region was set as equal to the smaller of the number of reads mapped to the forward strand or the number of reads mapped to the reverse strand.

After constructing the union set of each cell line, we summed the CP values of all sub-regions from the same sample to obtain the CP value for the merged region. The resulting CP values were used as read count inputs for DESeq2 analysis to detect differential bi-directional transcription across experimental conditions within the same cell line. A count matrix was constructed with columns representing the samples and rows representing the merged regions. To reduce the number of fragmented regions, we excluded regions with fewer than 5 CP in at least half of the samples. DESeq2 was run with default parameters, and regions with a *p_adj_* value less than 0.1 and a log2 fold change with an absolute value greater than 1 were considered as significant.

### Quantitation and statistical analysis

Statistical analysis of ATP bioluminescence cell viability assays, stress granule quantitation, and total PKR quantitation was performed with Prism version 7 (GraphPad). Statistical analysis of large datasets obtained from the Cancer Dependency Map or bulk RNA-sequencing data was performed with RStudio. The statistical tests used for each displayed graph is specified in the corresponding figure legends.

